# Neurodegenerative fluid biomarkers are enriched in human cervical lymph nodes

**DOI:** 10.1101/2024.04.26.591238

**Authors:** Adam Al-Diwani, Nicholas M Provine, Andrew Murchison, Rhiannon Laban, Owen J Swann, Ivan Koychev, Fintan Sheerin, Sandro Da Mesquita, Amanda Heslegrave, Henrik Zetterberg, Paul Klenerman, Sarosh R Irani

**Affiliations:** Department of Psychiatry, University of Oxford, Oxford, UK; Pandemic Sciences Institute, Nuffield Department of Medicine, University of Oxford, Oxford, UK; Department of Radiology, Oxford University Hospitals, Oxford, UK; UK Dementia Research Institute, Fluid Biomarker Laboratory, London, UK; Department of Neuroscience, Mayo Clinic, Jacksonville, Florida, USA; Department of Neurodegenerative Disease, UCL Institute of Neurology, Queen Square, London, UK; Department of Psychiatry and Neurochemistry, Institute of Neuroscience and Physiology, the Sahlgrenska Academy at the University of Gothenburg, Mölndal, Sweden; Clinical Neurochemistry Laboratory, Sahlgrenska University Hospital, Mölndal, Sweden; Hong Kong Center for Neurodegenerative Diseases, Clear Water Bay, Hong Kong, China; Wisconsin Alzheimer’s Disease Research Center, University of Wisconsin School of Medicine and Public Health, University of Wisconsin-Madison, Madison, WI, USA; Translational Gastroenterology Unit, Nuffield Department of Medicine, University of Oxford, Oxford, UK; Peter Medawar Building for Pathogen Research, Nuffield Department of Medicine, University of Oxford, Oxford, UK; Oxford Autoimmune Neurology Group, Nuffield Department of Clinical Neurosciences, University of Oxford, Oxford, UK; Department of Neurology, Mayo Clinic, Jacksonville, Florida, USA

## Abstract

In animal models, brain neurodegeneration biomarkers drain into cervical lymph nodes (CLNs). If this occurred in humans, CLNs may provide a readily accessible source of these biomarkers, draining the site of primary pathology. We tested this hypothesis in discovery and validation cohorts using ultrasound-guided fine needle aspiration (FNA).

We measured amyloid-beta 40 and 42, phospho-Tau-181, glial-fibrillary-acidic-protein, and neurofilament-light using single molecule array in CLN aspirates and plasma from: i) a discovery cohort of 25 autoimmune patients, and from ii) plasma, CLNs and capillary blood in four healthy volunteers, an optimisation-validation cohort.

FNA was well-tolerated by all participants. In both cohorts, all biomarkers were detected in all plasmas and CLNs, other than neurofilament-light (8/17 of discovery cohort). CLN biomarker concentrations were significantly greater than plasma concentrations for all except neurofilament-light, most markedly for phospho-Tau-181 (266 fold; *P*<0.02), whose CLN concentrations decreased with age (Spearman *r*=-0.66, *P*=0.001).

This study presents the first evidence that neurodegenerative biomarkers are detectable in human CLNs. Raised CLN:plasma biomarker ratios suggest their concentration in CLNs, which may offer a sensitive compartment for minimally-invasive sampling in clinical trials. Further, age-associated phospho-Tau-181 reduction with age suggests FNA of CLNs may measure the integrity of brain lymphatic drainage *in vivo*.

## Introduction

Neurodegenerative conditions are characterised by the accumulation of abnormal proteins in the brain. The pathological hallmarks of Alzheimer’s disease are amyloid-beta (Aβ) plaques and hyper-phosphorylated tau tangles.^1^ These, and other related neurodegenerative biomarkers, can be measured in cerebrospinal fluid (CSF) and blood, offering valuable insights into pathophysiology as well as clinical diagnosis, prognosis, and monitoring of Alzheimer’s disease and other dementias.^2^

Understanding the cross-compartmental dynamics of these proteins may be key to assessing their impaired clearance from the brain in dementias. Efforts largely focused in animal models, have recently discovered glymphatic and meningeal lymphatic vascular systems as candidates for extracellular fluid and macromolecule efflux from the brain into the subarachnoid space.^3–6^ Tracking CSF-injected molecules in mice, and few intrathecal gadolinium-based contrast studies in humans, suggest this brain meningeal lymphatic outflow drains directly into the cervical lymph nodes (CLNs).^5, 7–13^ Indeed, disruption of meningeal lymphatic drainage into the CLNs in mice can accelerate brain Aβ plaque formation and even alter its clearance with Aβ-immunotherapy.^5, 6, 14^ Despite this clear theoretical rationale, neurodegenerative biomarkers have yet to be measured in human CLN fluid-phase samples.

Previously, we have safely accessed CLNs in human participants using ultrasound-guided fine needle aspiration (FNA). These FNAs successfully identified both CNS antigen-directed lymphocytes and an abundance of neural proteins^15–17^: both show the enrichment of brain-specific markers within CLNs. Here, we significantly extend these concepts to test the hypothesis that fluid biomarkers relevant to the neurodegenerative processes are quantifiable in the CLNs and enriched compared to blood. If so, sampling from this anatomical site has the potential to illuminate both brain physiology and to complement CSF and blood analyses in neurodegeneration trials.

## Materials and methods

### Participants

A discovery cohort consisted of previously acquired frozen CLN aspirates and matched plasma samples from 25 participants with autoimmune neurological diseases (16 CLN aspirates and 23 plasma samples; median age 60 years, range=22-84, 14 female; Supplementary Table 1).^15,16^ To study fresh samples and exclude potential disease and sampling method-related confounding, we prospectively recruited four healthy adult volunteers with no self-reported or formal diagnoses of cognitive impairment (optimisation-validation cohort: mean age 33 years, range 24-38; three male; Supplementary Table 2). All provided informed written consent to donate CLN aspirates and paired peripheral blood (venous and capillary), in accordance with the Declaration of Helsinki and ethical approval (Research Ethics Committee 16/YH/0013).

### Sampling

#### Cervical lymph node fine needle aspiration

All FNAs were performed by a senior radiologist in a clinical ultrasound suite, as previously described.^15–17^ In brief, CLNs were visualised under ultrasound guidance and, after skin sterilisation, multiple passes were performed on the same node using a 23G hypodermic needle. Approximately 30μl of material was obtained and immediately after each needle withdrawal ice-cold sterile phosphate buffered saline (PBS) solution was aspirated through the needle bevel into a syringe and the material then ejected into a 1.5ml microtube. The discovery cohort focussed on cell recovery rather than supernatant optimisation such that needle passes were repeated 2-3 times with varying volumes (0.5-2ml) for each needle wash. These were pooled resulting in variable total supernatant volumes (1.5-8ml). In the optimisation-validation cohort, we standardised supernatant acquisition with exactly two needle passes, and material was carefully washed into a separate 1.5ml microtube for each of four washes per needle. In both cohorts, following transfer to the laboratory on ice, samples were centrifuged for five minutes (200G-400G), and supernatants stored at −80°C.

#### Capillary blood

Capillary blood was also collected by lancet in order to sample a micro-vascularised tissue akin to the lymph node hilum which lacked a specialised lymphatic confluence. One drop (∼30μl) was collected into a 1.5ml microtube and, as per the lymph node FNA procedure, aspirated into a 23G hypodermic needle followed by a 1ml of ice-cold PBS pull-through. Thereafter, the needle was removed and the mixture ejected into a fresh 1.5ml microtube. The diluted capillary blood was centrifuged at 400G for 5 min. The acellular supernatant was transferred to a fresh 1.5ml microtube, and stored at −80°C. The tube lacked anti-coagulant but there was no visible clotting. We refer to the diluted capillary plasma as capillary blood supernatant.

#### Venous blood

Standard venepuncture was also conducted to collect venous blood in an EDTA tube for plasma, as used here, and for PBMC isolation reported previously.^17^ Following transfer to the laboratory on ice, blood was layered over 15ml lymphoprep and centrifuged at 931G for 30 minutes with no brake. Plasma was transferred to 1.5ml microtubes, and stored at −80°C.

#### Assays

The samples were assayed on the ultrasensitive Quanterix HD-X Single molecule array (SIMOA) platform: a Human Neurology 4-Plex E (N4PE) kit for two isoforms of amyloid-beta (Aβ)_40_ and _42_, glial fibrillary acidic protein (GFAP) and neurofilament light (NfL), and a monoplex phosphorylated tau 181 Advantage V2.1 kit (pTau181). To optimise assays of CLN and capillary supernatants, serial dilutions in sample buffer found a 2-fold dilution to be optimal. Raw concentrations were multiplied by this assay dilution factor. Each sample was run in duplicate and the mean reported. Concentrations were further multiplied by a correction factor to account for dilution at the time of sampling (dilution volume/material volume, which for the validation cohort was 1000μl/30μl = 33.33).

#### Statistical analysis

Analyses were performed in Excel v16.83 (Microsoft), and Prism v10.2.1 (GraphPad). Data were rounded to three significant figures to avoid pseudo-exactness given known immunoassay coefficient of variation (5-10%). Paired data were compared using a Wilcoxon signed-rank test or a two-tailed paired Student’s *t-test* for non-normal and normally distributed data, respectively. Unpaired non-normally distributed data were compared using a two-tailed Mann-Whitney test. Correlation was estimated with Spearman’s rank co-efficient. Significance is indicated by asterisks: \**P* ≤ 0.05, \*\**P* ≤ 0.01, \*\*\**P* ≤ 0.001.

## Results

All participants underwent sampling without any physical or psychological adverse events. In the discovery cohort, all five biomarkers were detected in all of the CLN samples, with the exception of NfL which was detectable in 8/17 (Fig. 1A-B). The corrected CLN concentrations of Aβ_40_, Aβ_42_ and pTau181 were significantly higher than those in plasma (*P*=0.03, 0.005, and <0.001, respectively, two-tailed Wilcoxon signed-rank test). For pTau181, the magnitude of difference was greatest, ∼65-fold with a median CLN corrected concentration of 1465 pg/ml (range 176-18900) versus plasma median 22.6 pg/ml (range 13.3-43.6). No significant differences were noted for GFAP and NfL.

**Figure 1.**
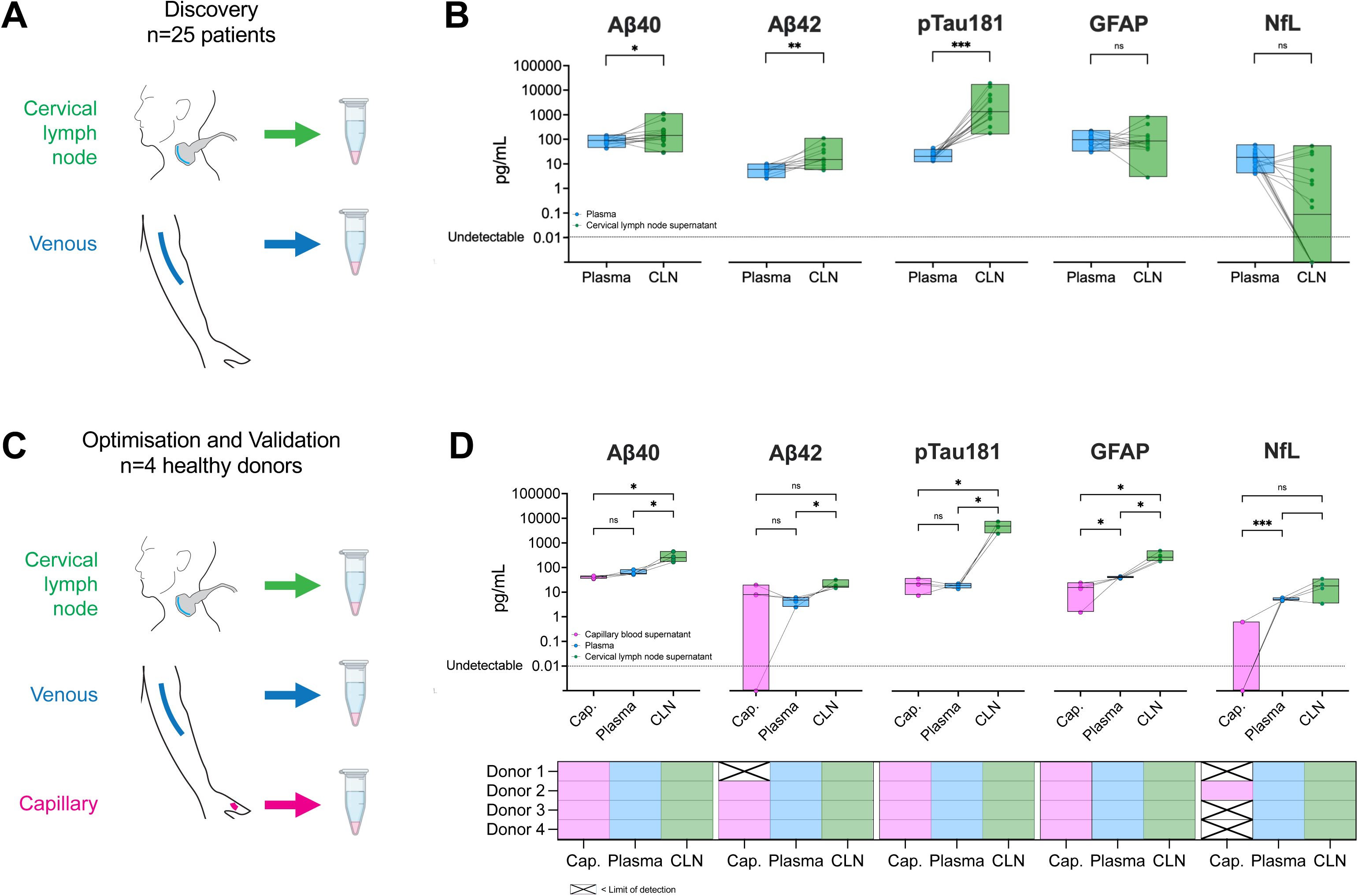
Corrected concentrations of dementia fluid biomarkers in plasma versus capillary and cervical lymph node supernatants. (A) Schematic diagram illustrates sample sources (venous blood as plasma and cervical lymph node aspirate supernatant) from 25 clinical donors with autoimmune neurological diseases which acted as a discovery cohort. (B) Dot and box plots indicate the concentrations of five dementia fluid biomarkers from two colour-coded bio-samples from 25 individuals on a logarithmic axis (plasma = blue, cervical lymph node supernatant = green). The box spans the minimum to maximum value with a horizontal line at the median. Lines between dots indicate the same individual across the two sample types. A dotted line runs across the graphs to indicate the limit of assay detection. The result of a Wilcoxon signed-rank test for each protein is indicated above the plots. (C) Schematic diagram illustrates sample sources (venous blood as plasma, capillary bed blood as supernatant, and cervical lymph node aspirate supernatant) from four healthy donors which acted as an optimisation-validation cohort. (D) Dot and box plots indicate the concentrations of five dementia fluid biomarkers from three colour-coded bio-samples from four individuals on a logarithmic axis (capillary blood supernatant = pink, plasma = blue, cervical lymph node supernatant = green). The box spans the minimum to maximum value with a horizontal line at the median. Lines between dots indicate the same individual across each sample type. A dotted line runs across the graphs to indicate the limit of assay detection. The result of a two-tailed paired *t-test* for each protein is indicated above the plots. The results for the above protein are summarised below for each donor and sample type. Cells are filled with the same colour scheme to indicate a positive result and a X indicates that this was below the limit of detection. Abbreviations used: Aβ = amyloid-beta peptide; Cap. = capillary blood supernatant; CLN = cervical lymph node supernatant; GFAP = glial fibrillary acidic protein; NfL = Neurofilament light; ns = not significant; pg/mL= picogram per millilitre; pTau181 = phosphorylated tau protein 181. \**P* ≤ 0.05, \*\**P* ≤ 0.01, \*\*\**P* ≤ 0.001.

In the optimisation-validation cohort, we observed that the second FNA needle pass yielded more tightly distributed concentrations of these biomarkers, consistent with our previous proteomic observations (Supplementary Fig. 1A).^17^ More specifically, the first wash from this second needle consistently showed the highest concentration and tightest grouping (Supplementary Fig. 1B). Using this sample, all biomarkers were detectable in all CLN supernatants and paired plasma in all four participants (Fig. 1C-D). In contrast, Aβ_42_ and NfL were undetectable in one and three of the four capillary supernatants, respectively. All CLN samples showed higher corrected concentrations of Aβ_40_, Aβ_42_, pTau181, and GFAP compared to plasma (*P*=0.04, 0.05, 0.02, and 0.03, respectively, two-tailed paired *t-test*; Fig. 1D), and of Aβ_40_, pTau181 and GFAP compared to capillary supernatants for (*P*=0.03, 0.02, and 0.03, respectively, two-tailed paired *t-test*; Fig. 1D). There were no statistically significant differences between plasma and capillary blood for Aβ_40_, Aβ_42_ and pTau181, although GFAP and NfL were significantly lower in the capillary samples (*P*=0.02 and 0.001, respectively, two-tailed paired *t-test*; Fig. 1D).

Consistent with the discovery cohort, the highest fold differences in corrected concentrations were between CLN compared to both plasma and capillary samples for pTau181: 266 times higher in CLN than plasma (mean 4865 vs. 18.3 pg/ml; *P*=0.02, two-tailed paired *t-test*), without differences between capillary and blood plasma (18.4 vs 21.8 pg/ml, ratio=0.84; *P*=0.65, two-tailed paired *t-test*; Fig. 1D). While the absolute mean corrected concentration of CLN Aβ_40_ was higher than Aβ_42,_ the ratio of each versus their respective plasma samples was similar (4.5) and the Aβ_42/40_ ratio, a commonly used dementia biomarker index, was not different between plasma and CLN (Supplementary Fig. 2).

To explore the hypothesis that lymphatic drainage reduces with age, we combined both groups to yield a broad age range (*n*=29; median 56 years, range 22-84). In this exploratory analysis while, as could be expected, plasma levels positively correlated with age (Spearman *r*=0.59, *P*=0.001), CLN levels showed a negative correlation of pTau181 versus age (Spearman *r*=-0.65, *P*=0.002) (Fig. 2). There was no correlation between CLN and age for the other four biomarkers, but the levels of GFAP and NfL in the plasma positively correlated with age (Spearman *r*=0.69, *P*<0.0001 and *r*=0.81, *P*<0.0001, respectively; Supplementary Fig. 3).

**Figure 2.**
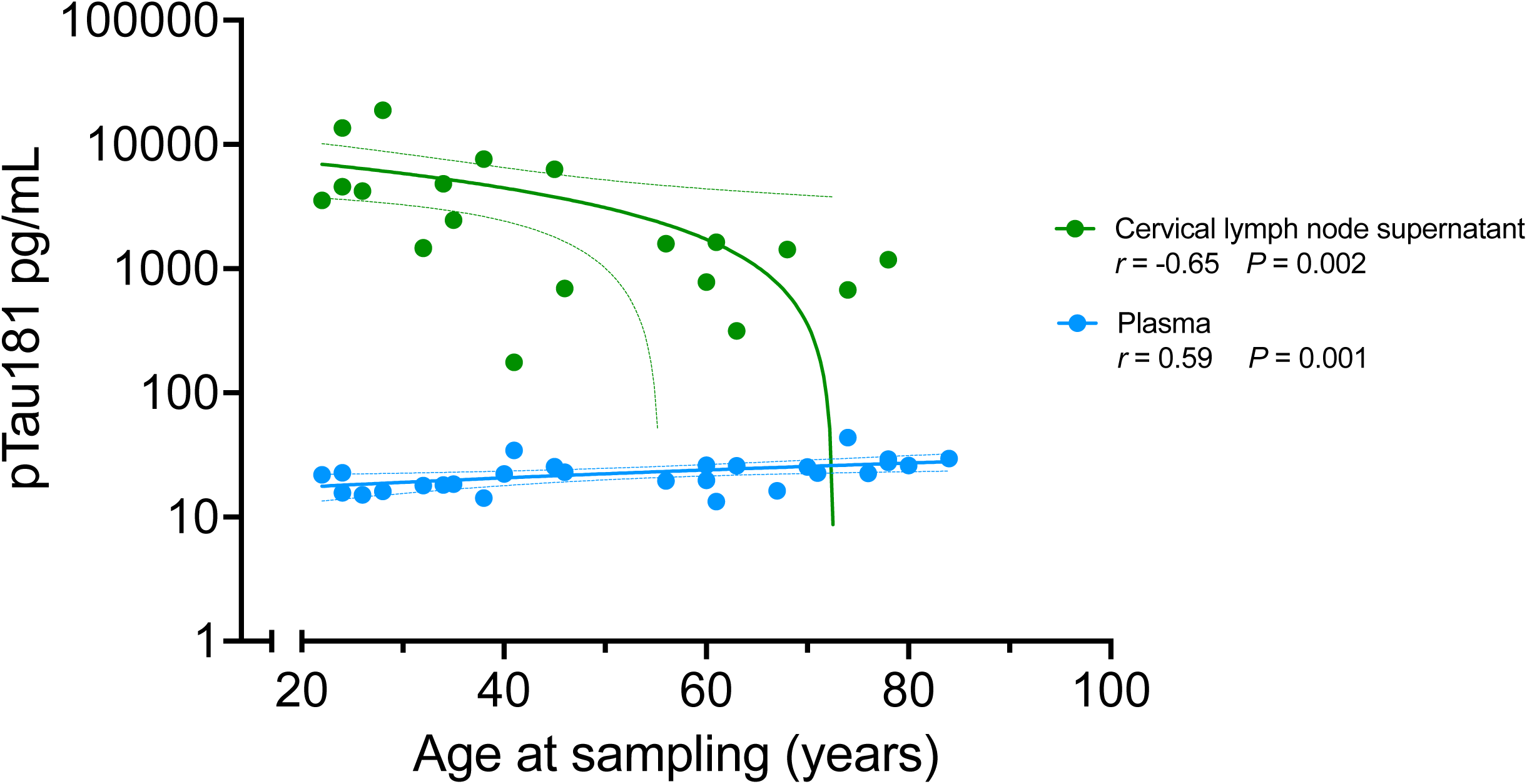
Association of phosphorylated tau 181 (pTau181) concentration in cervical lymph node and plasma with age. Correlation of corrected concentration of pTau181 in plasma and cervical lymph node supernatant is plotted against age (cervical lymph node=green, plasma=blue). Each dot is one sample and a line of best fit is drawn to summarise the relationship with dotted lines indicating the 95% confidence interval. The Spearman correlation co-efficient (*r*) and statistical significance (*P*) for each line is given above the graph. Abbreviations used: CLN = cervical lymph node, pg/mL= picogram per millilitre, pTau181 = phosphorylated tau 181

## Discussion

Our results show that CLN aspirates consistently contain and concentrate proteins that are currently used as neurodegenerative biomarkers. Given the direct and intimate connections between the brain and CLNs, mediated by meningeal lymphatics, we propose CLNs may represent a highly sensitive source of material, which is safe and minimally invasive to sample. Hence, we propose cross-sectional and longitudinal sampling of CLNs may be as part of future biological and clinical studies in neurodegeneration.

In the technically standardised young adult validation group, CLN biomarker levels were tightly-grouped and showed higher concentrations than blood. This suggests these proteins are being physiologically concentrated in CLNs, rather than reflecting blood contamination. This anatomical compartmentalisation is consistent with clear distinctions in cellular populations we have previously observed when sampling paired CLNs and blood, such as enrichment for lymph node resident T follicular helper cells.^15–17^

The next critical question is whether these proteins are being enriched in CLNs via brain lymphatic drainage, non-brain head and neck drainage, local production, or a combination. One clue comes from the relative neural specificity of the proteins described here. In our prior study gene ontology commonly ascribed neural functions to CLN-restricted proteins. Examples included secernin-1, a tau-binding protein^18^, the key microglial receptor TREM2^19^, and ADAM22, a key receptor of the neuronal autoantibody target LGI1.^20^ It may also be that local lymph node sources are relevant. For example, TREM2 itself is expressed by lymph node sub-capsular macrophages^21^ and both Aβ and phosphorylated tau can be found in the periphery including the head and neck.^22–24^ Nonetheless, as exemplified by a study of post-stroke tonsillar biopsies, these regions themselves can be differentially enriched for drained brain-derived proteins.^25^ Moreover, as several independent animal models have shown, both dynamically and at post-mortem, the CLNs receive CSF brain drainage from the meningeal lymphatics.^8–10^ Future studies should include concomitant disease biomarker measurements in CSF samples. We speculate that biomarker concentrations detected in the CLNs will better reflect brain CSF levels than plasma, due to the high and constant rate of CSF macromolecule drainage into the CLNs via meningeal lymphatics.^4–6, 13^

Presuming the CLN biomarkers are brain-derived, then the varied profiles merit consideration. In particular, the three log10-fold concentration difference of pTau181 in CLN versus blood is striking. pTau181 in blood and CSF is associated with the clinical phase of Alzheimer’s disease and cerebral atrophy^26^, yet the validation group donors were young and with no self-reported or formal diagnoses of cognitive impairment. This could indicate a constitutive drainage of phosphorylated tau isoforms by the meningeal lymphatic system. Their build up in the extracellular compartment in older people, possibly due to decreased drainage into the CLNs, may therefore contribute to the prion-like tau propagation seen during disease progression in primary and secondary tauopathies.^27^ The reduction in CLN pTau181 levels with age identified in our initial exploratory analysis could fit with this brain lymphatic drainage model.

Yet, despite their detection at higher concentrations in the CLNs, neither Aβ40 and Aβ42 had such a similar magnitude difference between the two sites nor did the CLN concentration correlate with age. The amyloid-beta precursor protein is widely expressed across extra-cerebral tissues, including blood platelets, myocytes, and hepatocytes, from which both Aβ40 and Aβ42 are proteolytically produced and released.^28^ Aβ peptides have a high propensity for extracellular aggregation in the form of plaques, which form at a faster rate both in the brain and meninges in ageing Alzheimer’s disease transgenic mice.^29^ The aggregation propensity of Aβ might preclude its presence in the CSF that reaches the dura for drainage by the meningeal lymphatic vessels. It may also be that Aβ peptides molecules are cleared by a cellular route. Indeed, a histological study of pathological lymph nodes showed Aβ-staining cells were considerably more common in the cervical than the inguinal lymph nodes.^30^ Moreover, it may be that only a certain proportion of specific brain-derived proteins drain from meningeal lymphatics into the CLNs and so certain proteins may show alternative patterns. In contrast, the lack of difference for NfL is more explicable since elevated extracellular NfL demarcates neuro-axonal damage usually found in aggressive neuro-degenerative or neuro-inflammatory processes. Thus, the lack of elevated NfL in the CLNs is consistent with the profile of the donors.

Limitations include the small numbers in our validation cohort and potential confounds including use of a variety of autoimmune neurological conditions and differences in sample preparation. However, even despite these aspects the main observations are consistent across the two independent cohorts, and the technically optimised CLN data are tightly grouped in our four healthy individuals. Further, from a technical perspective, we highlight the differential abundance of these biomarker proteins between an initial and second needle pass – consistent with our observations at proteomic scale.^17^ We propose this reflects a degree of material dissociation in CLNs which is then more easily aspirated following the first needle pass. This observation will help future groups perform the highest yield aspirations. Ultimately, larger cohort sizes but also complementary data including lymph node sampling distant from the head and neck, CSF, imaging indices, as well as both steady state and natural history, experimental, or clinical interventions will be needed to further elucidate. Nonetheless, the large effect sizes seen in this pilot study together with our previous immunologic and proteomic data suggests we are tapping into a wealth of brain-relevant material in human CLNs.

In conclusion, we provide independently validated evidence to support the presence of dementia fluid biomarkers in CLNs, in all except NfL, at significantly higher levels than circulating plasma. Moreover, we provide the first evidence in humans of reduced brain lymphatic drainage of the disease-relevant biomarker pTau181 with age. Taken together, these observations likely reflect potential to longitudinally quantify *in vivo* lymphatic drainage of brain disease biomarkers from CLN aspirates. Hence, this minimally invasive technique is likely of great value for human experimental medicine studies and early phase clinical trials. Given the relative ease and excellent safety profile of CLN FNA, we propose the technique is now ready to move from the rare autoimmune neurological diseases in which we pioneered its application, to healthy ageing, and more common brain disorders such as infection, traumatic brain injury, and dementia.

## Data availability

Data supporting the study are available from the corresponding authors upon reasonable request.

---

**Supplementary Figure 1.**
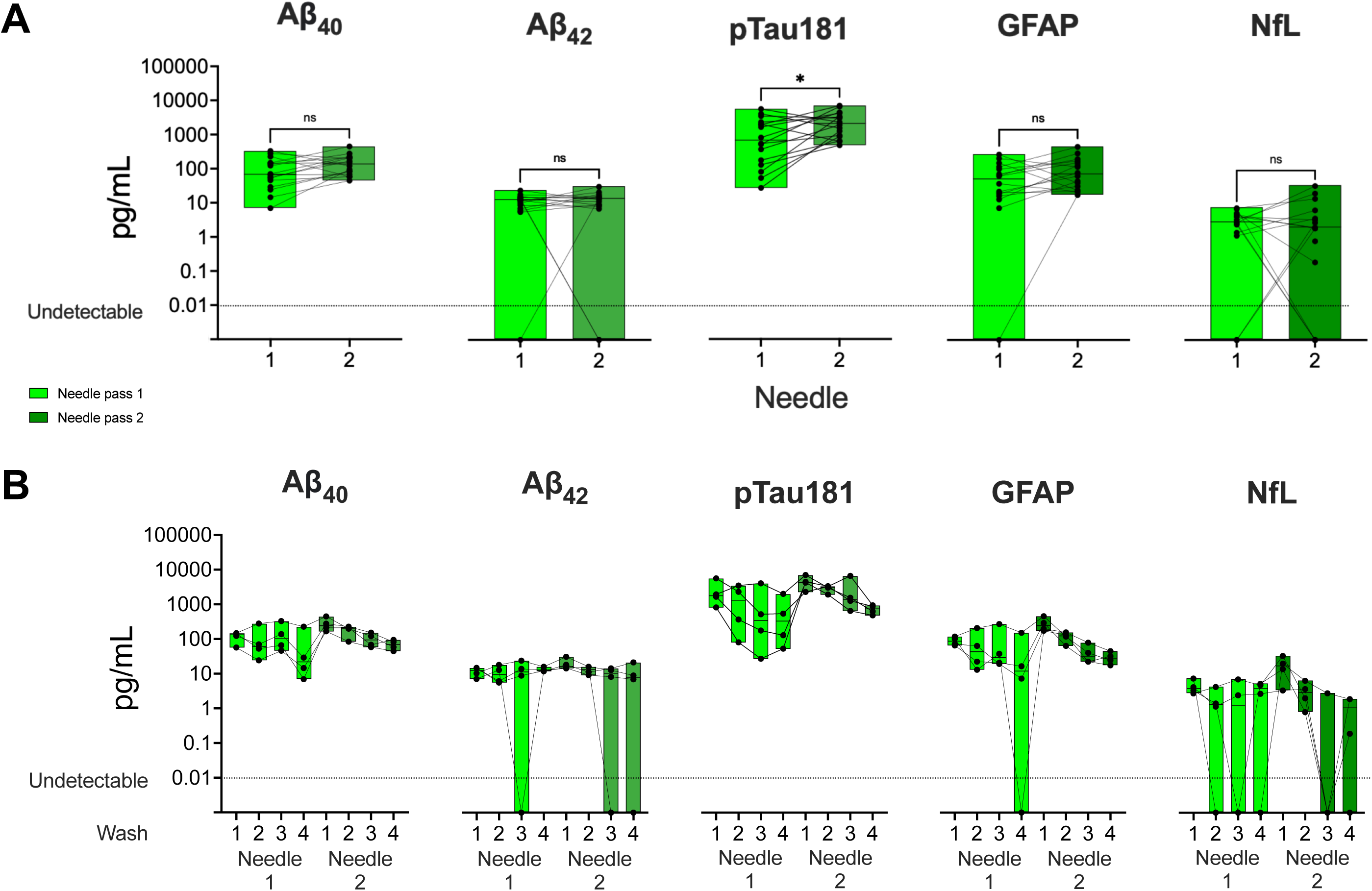

**Supplementary Figure 2.**
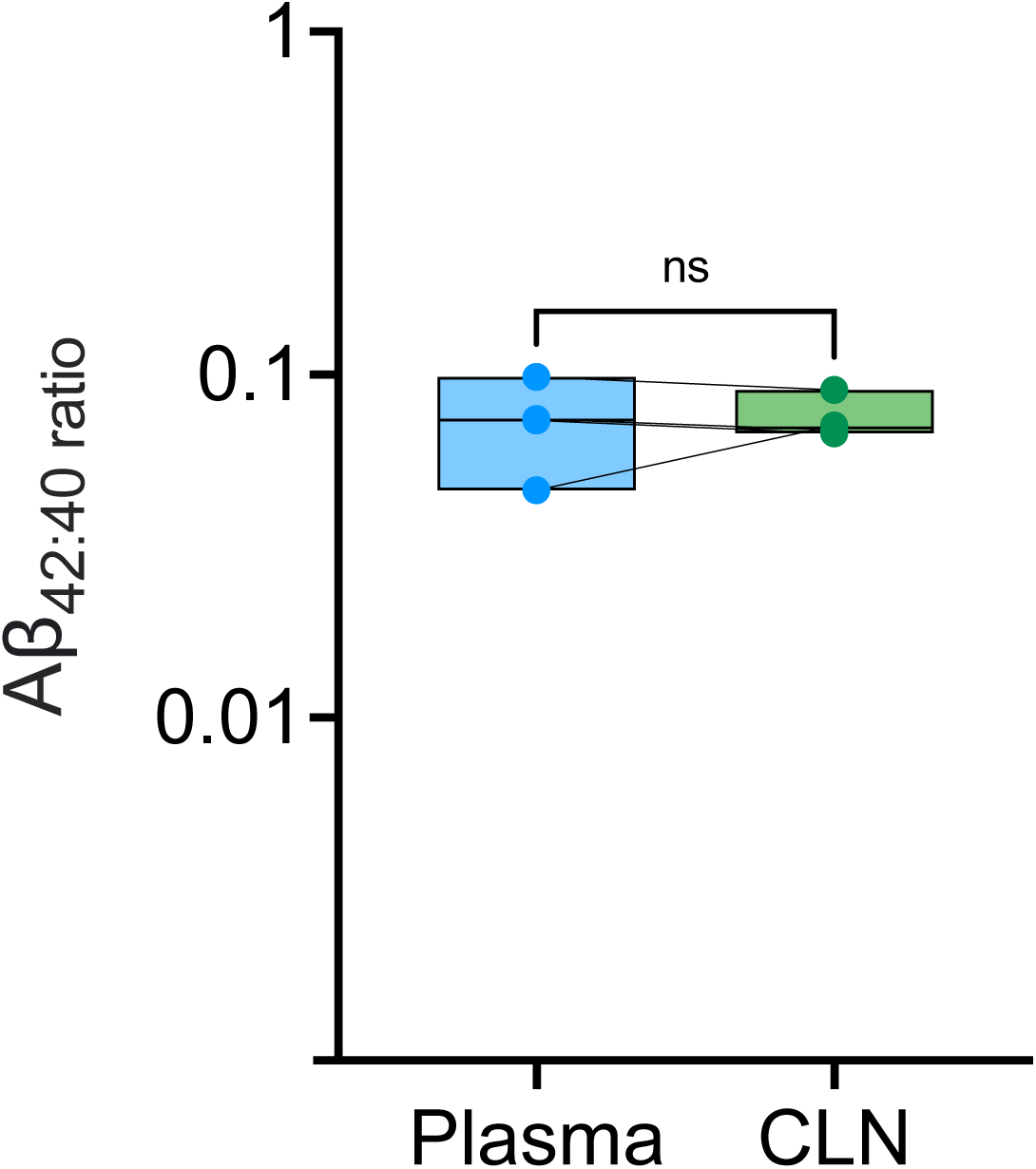

**Supplementary Figure 3.**
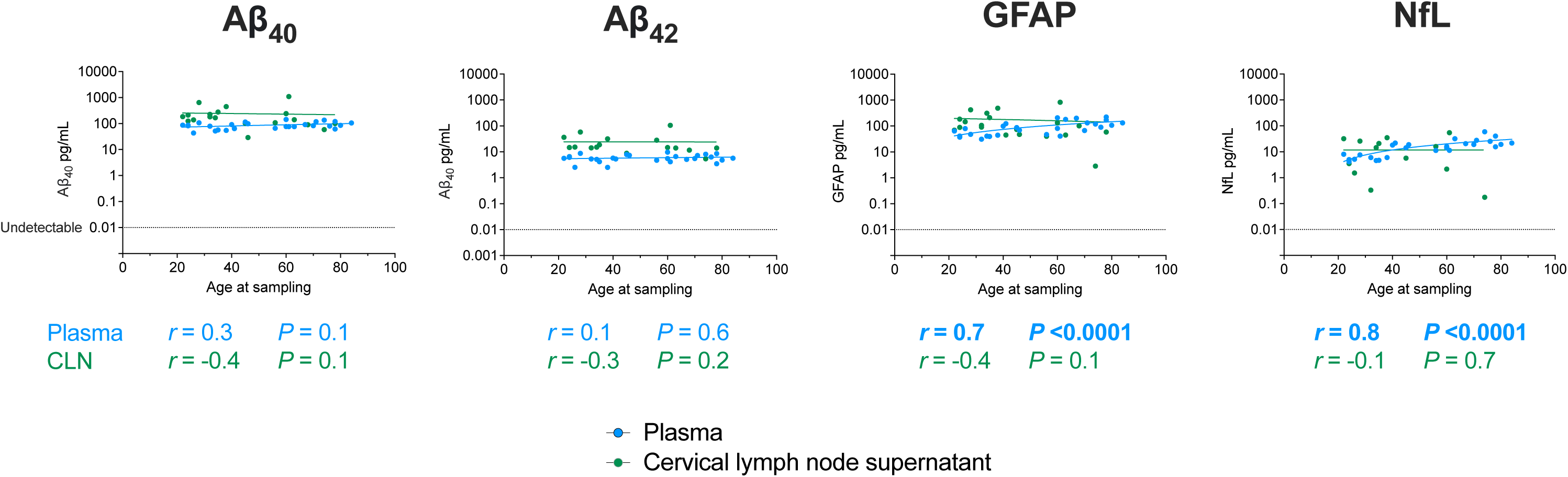

