## Supplementary material for "Neurodegenerative fluid biomarkers are enriched in human cervical lymph nodes"

**Supplementary figure 1 Corrected concentrations of dementia fluid biomarkers in cervical lymph node supernatants from four serial washes of two separate fine needle aspirate passes**

(A) In four individuals, dot and box plots indicate on a logarithmic axis the concentrations of five dementia fluid biomarkers from cervical lymph node fine needle aspirate from four consecutive washes of two consecutive fine needle passes of the same node (needle one = light green; needle two = dark green). The data are pooled into needle 1 and 2 and compared using a Mann-Whitney test. A dotted line indicates the limit of detection.

(B) The same data are presented on a per wash basis plotted consecutively from left to right with wash and needle numbers annotated. Lines between dots indicate the same individual across each needle-wash sample.

Abbreviations used:

Aβ = amyloid-beta peptide; GFAP = glial fibrillary acidic protein; NfL = Neurofilament light; pg/mL= picogram per millilitre; ns = not significant, pTau181 = phosphorylated tau 181 protein. *$P \leq 0.05$.

**Supplementary Figure 2 Amyloid-beta 42 to 40 ratio in plasma versus cervical lymph node aspirate supernatant samples**

Dot and box plots indicate on a logarithmic axis indicate the ratios of amyloid-beta peptides 42 and 40 in plasma and cervical lymph node aspirate samples in four individuals (plasma = blue, cervical lymph node supernatant = green). The box spans the minimum to maximum value with a horizontal line at the median. Lines are drawn to indicate the same individual across the two sample types. The result of a paired t-test comparison is indicated above the plot.

Abbreviations used:

Aβ = amyloid-beta peptide; CLN = cervical lymph node supernatant; ns = not significant; pg/mL= picogram per millilitre

**Supplementary Figure 3 Association of dementia biomarker concentrations in cervical lymph node and plasma with age**

Biomarker concentrations are plotted against age for plasma and cervical lymph node aspirate supernatant samples (A$\beta_{40,42}$, GFAP, and NfL; plasma = blue, cervical lymph node supernatant = green). Each dot is one sample and a line of best fit summarises the relationship. A dotted line indicates the limit of assay detection. The Spearman correlation co-efficient ($r$) and statistical significance ($P$) for each line is given below each graph rounded to one significant figure. Statistically significant results are indicated in bold.

Abbreviations used:

Aβ = amyloid-beta peptide; CLN = cervical lymph node, GFAP = glial fibrillary acidic protein, NfL = Neurofilament light; ns = not significant, pg/mL= picogram per millilitre.

**Supplementary Table 1 Clinical and demographic details of additional cervical lymph node aspirate and plasms donors**

| Clinical | Age at sampling | Sex | Plasma sample available | CLN sample available |
|---|---|---|---|---|
| LGI1 | 40 | M | Yes | No |
| LGI1 | 60 | M | Yes | No |
| LGI1 | 67 | F | Yes | No |
| LGI1 | 70 | M | Yes | No |
| LGI1 | 71 | M | Yes | No |
| LGI1 | 74 | M | Yes | Yes |
| LGI1 | 76 | M | Yes | No |
| LGI1 | 78 | M | Yes | Yes |
| LGI1 | 78 | M | Yes | No |
| LGI1 | 80 | M | Yes | No |
| LGI1 | 84 | M | Yes | No |
| NMDAR | 28 | F | Yes | Yes |
| NMDAR | 22 | F | Yes | Yes |
| NMDAR | 26 | F | Yes | Yes |
| NMDAR | 32 | F | Yes | Yes |
| NMDAR | 32 | F | No | Yes |
| NMOSD | 24 | F | Yes | Yes |
| NMOSD | 41 | F | Yes | Yes |
| NMOSD | 45 | F | Yes | Yes |
| NMOSD | 46 | F | Yes | Yes |
| NMOSD | 56 | M | Yes | Yes |
| NMOSD | 60 | F | Yes | Yes |
| NMOSD | 63 | F | Yes | Yes |
| GlyR | 61 | F | Yes | Yes |
| GAD | 68 | F | No | Yes |

25 donors with five different autoimmune neurological disorders donated paired cervical lymph node and blood samples, of which 16 aspirates and 23 plasma samples were available to analyse for this study.

Abbreviations used:

CLN = cervical lymph node, GAD = Glutamic acid decarboxylase antibody-associated disorder, GlyR = Glycine receptor-antibody associated disorder, LGI1 = Leucine-rich-glioma-inactivated 1-antibody encephalitis, NMDAR = N-methyl-D-aspartate receptor-antibody encephalitis, NMOSD = Neuromyelitis optica spectrum disorder

**Supplementary Table 2 Corrected concentrations of dementia fluid biomarkers in optimisation cohort plasma versus capillary and cervical lymph node supernatants**

| Sample properties | | | | Corrected mean concentration (pg/mL) | | | | |
|---|---|---|---|---|---|---|---|---|
| Donor | Age at sampling (Years) | Sex | Material | Aβ40 | Aβ42 | pTau181 | GFAP | NfL |
| 1 | 38 | M | Capillary | 34.5 | <LOD | 21.3 | 1.6 | <LOD |
| | | | Plasma | 54.5 | 2.5 | 14.2 | 44.2 | 6 |
| | | | CLN (Va) | 447 | 31.8 | 7610 | 487 | 34.9 |
| 2 | 34 | F | Capillary | 44.7 | 8 | 22.2 | 16.4 | 0.6 |
| | | | Plasma | 51.9 | 5.1 | 18.1 | 40.7 | 4.6 |
| | | | CLN (Va) | 167 | 15.1 | 4810 | 311 | 14.5 |
| 3 | 24 | M | Capillary | 36.7 | 7.7 | 7.6 | 24.2 | <LOD |
| | | | Plasma | 82.2 | 6.1 | 22.7 | 37.2 | 5 |
| | | | CLN (Va) | 213 | 14.7 | 4580 | 184 | 3.5 |
| 4 | 35 | M | Capillary | 45.5 | 19.6 | 36.1 | 14.2 | <LOD |
| | | | Plasma | 56.9 | 4.2 | 18.4 | 39.4 | 4.7 |
| | | | CLN (Ia) | 277 | 18.7 | 2460 | 211 | 20.9 |
| Mean (SD) | 33 (6) | - | Capillary | 40.4 (5.5) | 8.8 (8) | 21.8 (11.6) | 14.1 (9.4) | 0.2 (0.3) |
| | | | Plasma | 61.4 (14) | 4.5 (1.5) | 18.4 (3.5) | 40.4 (2.9) | 5.1 (0.6) |
| | | | CLN | 276 (123) | 20.1 (8) | 4865 (2114) | 298 (137) | 18.5 (13.1) |

Abbreviations used:

Aβ = amyloid-beta peptide; Capillary = capillary blood supernatant; CLN = cervical lymph node supernatant Pass 2-Wash 1, anatomical level indicated in parentheses; F = female; GFAP = glial fibrillary acidic protein; M = male; <LOD less than limit of detection of assay; NfL = Neurofilament light; pg/mL= picogram per millilitre; pTau181 = phosphorylated tau 181 protein; SD = standard deviation
